# Genome-scale metabolic model atlas of the zoonotic pathogen *Streptococcus suis*

**DOI:** 10.64898/2026.06.17.732996

**Authors:** Karl Kochanowski, Chenxi Liu, Pau Obregon-Gutierrez, Gemma G. R. Murray, Muriel Dresen, Ines Lefranc, Heather Wells, Ayelén Perez-Falcón, John T. Munnoch, Paul A. Hoskisson, Daniel Machado, Alexander W. Tucker, Florencia Correa-Fiz, Virginia Aragon, Lucy A. Weinert

**Author notes:** Department of Veterinary Medicine, Institute of Microbiology and Epizootics, Freie Universität Berlin, Berlin, Germany (Muriel Dresen). **Shared-first authors**: Karl Kochanowski, Chenxi Liu. **Corresponding author**: Karl Kochanowski.

## Abstract

*Streptococcus suis* is a Gram-positive bacterium with a dual role as a commensal member of the porcine nasal microbiota and a pathogen causing systemic disease in pigs and humans. Mounting evidence suggests that metabolism is a key driver of *S. suis* pathogenicity. Given the species’ high genetic variability, we hypothesize that differences in metabolic networks could explain the diverse pathogenic phenotypes observed across different strains. To test this, we generated an atlas of over 3000 strain-specific and automatically curated genome-scale metabolic models that cover the breadth of pathogenic and commensal *S. suis* lineages. Using this model atlas, we performed the first species-level examination of metabolic traits in *S. suis*. Our simulations, supported by experimental validation, revealed three key insights. First, while metabolic traits are broadly conserved in *S. suis*, there are nevertheless lineage-dependent differences in amino acid auxotrophies and carbon utilization patterns that point towards distinct *in vivo* niches. Second, most strains are predicted to grow in different plausible *in vivo* environments regardless of their virulence phenotype, suggesting that metabolism is a weak barrier to systemic infection. Third, by systematically predicting reaction essentiality in more than 15 million reaction-strain-condition combinations, we identify a subset of 17 reactions, largely in nucleotide metabolism, that are conditionally essential *in vivo* and may serve as new targets for the development of new antimicrobials or vaccines. Overall, this study provides a valuable new resource for broadly examining *S. suis* metabolism and its role in pathogenicity.

## Introduction

*Streptococcus suis* is a Gram-positive bacterium with an intriguing dual lifestyle: On the one hand, it is an important member of the porcine airway microbiota commonly found in the nasal cavity (*1*, *2*) and tonsils (*3*) of healthy animals. On the other hand, it is also a major pathogen that causes systemic disease in pigs as well as humans (*4*). Given this impact on pig production and human health, many recent efforts have focused on elucidating potential drivers of *S. suis* pathogenicity.

Mounting evidence from numerous *in vitro* and *in vivo* studies suggests that *S. suis* metabolism plays a key role in its virulence and adaptation to different host niches, as discussed in detail in a recent review (*5*). First, recent gene expression studies have shown that carbohydrate availability affects the *in vitro* expression of virulence genes (*6*, *7*) and many metabolic *S. suis* genes are upregulated *in vivo* during systemic infection in pigs (*8*). Second, various gene deletion studies have shown that the deletion of many metabolic genes affects the ability of *S. suis* to become systemic (*9–18*). Finally, metabolic proteins also moonlight as direct virulence factors, such as adhering to host cells and binding plasminogen (*19–21*). These studies highlight the functional importance of metabolism for *S. suis* pathogenicity. However, there are ongoing discussions on which of the many postulated metabolic virulence factors in *S. suis* are truly important for its pathogenicity (*22*), and our current understanding of the relationship between metabolism and pathogenicity is limited to a handful of laboratory-adapted strains. This is unfortunate as genomic variability in *S. suis* is vast (*23*), with extensive variation in pathogenicity through the emergence of highly pathogenic lineages (*24*).

Here, we aim to address these questions by examining the metabolic traits of the various circulating *S. suis* lineages using genome-scale metabolic models (GSMMs). GSMMs have emerged as powerful computational tools for studying strain-specific differences in metabolic traits in various bacterial pathogens (*25*, *26*), including *Escherichia coli* (*27–29*), *Staphylococcus aureus* (*30*), *Salmonella* (*31*), *Mycoplasma* (*32*, *33*), and *Klebsiella* (*34*, *35*). However, to date only a handful of *S. suis* metabolic models exist (*36*, *37*), thus hampering the systematic comparison of metabolic traits across *S. suis* pathogenic and commensal strains.

To overcome this limitation, we generated an atlas of over 3000 strain-specific and curated *S. suis* metabolic models. This strain collection included various distinct pathogenic lineages that emerged in the 19^th^ and 20^th^ century and are strongly associated with disease in pigs and humans, numerous commensal lineages, and groups of isolates that are evolutionarily divergent from the central population of *S. suis* (i.e. out-groups) (*24*). Using this model atlas, we provide the first species-level examination of metabolic traits in *S. suis*, examining how nutrient requirements, the ability to grow in different plausible *in vivo* environments, and reaction essentiality differ across lineages. Our data reveal hallmarks of *S. suis* metabolism as well as numerous metabolic traits that diverge in selected lineages, and which we could experimentally validate *in vitro*. Overall, this study provides a valuable new resource for broadly examining *S. suis* metabolism and the role it plays in pathogenicity.

## Results

### Section 1: Generating an atlas of 3000+ strain-specific S. suis genome-scale metabolic models

Our goal was to generate a set of strain-specific metabolic models that would adequately represent the genetic variability of *S. suis* isolates. Towards this end, we leveraged a large collection of over 3000 fully sequenced *S. suis* isolates, which cover the full breadth of currently known pathogenic and commensal lineages (*24*). To generate strain-specific metabolic models for each of these isolates, we developed a scalable computational pipeline that combines automated model reconstruction (using CarveMe (*36*)) and additional curation based on currently available *in vitro* data (**Figure 1A**, see **methods** for details on the exact model reconstruction and curation steps we implemented). This pipeline enabled the reconstruction and automated curation of 3070 strain-specific models with high overall quality (MEMOTE (*38*) scores of > 70% in all cases, in line with current state of the art models, see **Data set 1**) in about 24h on a standard laptop.

**Figure 1.**
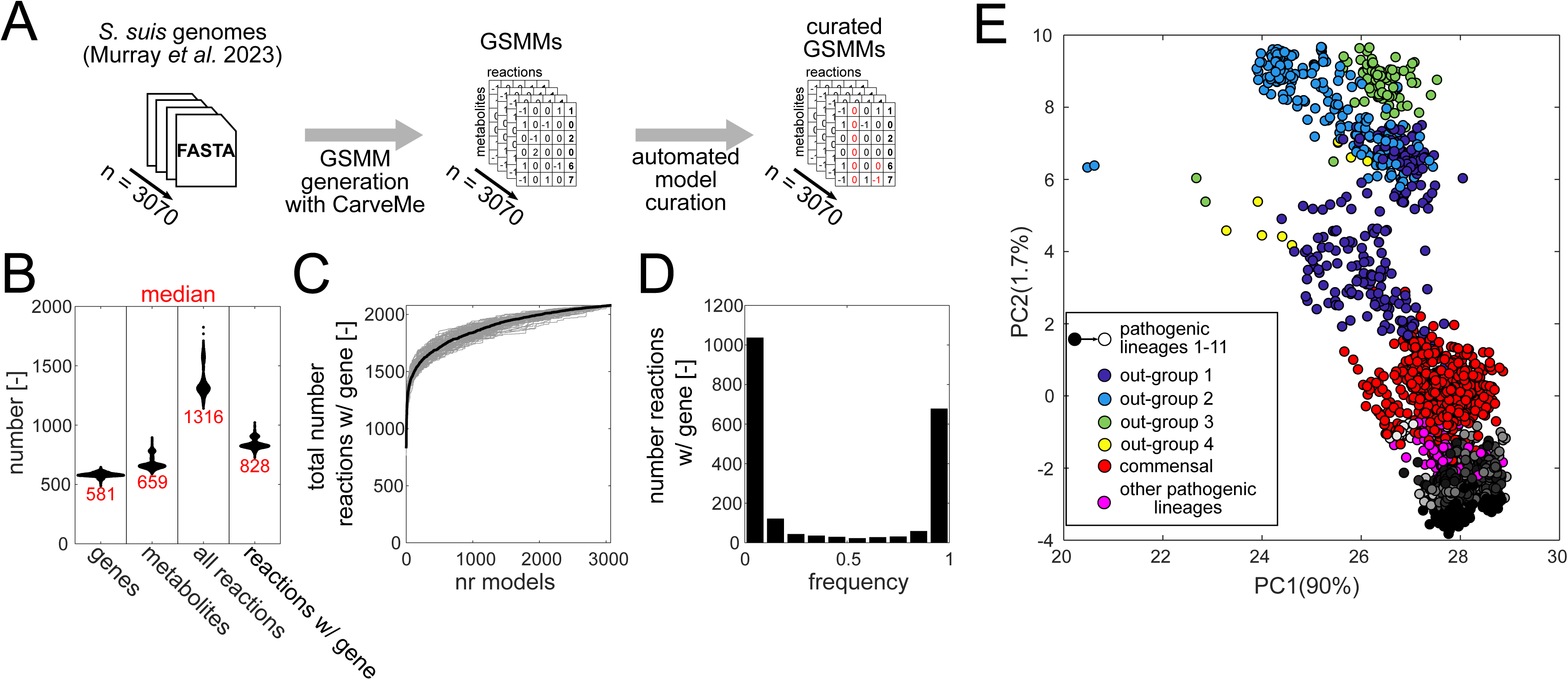
Atlas of 3070 strain-specific *Streptococcus suis* genome-scale metabolic models (GSMMs). **A)** Schematic of approach. **B)** Swarm chart showing distribution of (from left to right) the number of genes, the number of unique metabolites, the total number of reactions, and the number of reactions having a gene associated, across all models. Numbers in red: median values. **C)** The total number of gene-associated reactions as a function of the number of models considered. Gray lines: 100 iterations with randomized order in which the models were included. Black line: median. **D)** Distribution of frequency at which each gene-associated reaction is found across all models. **E)** Principal component analysis (PCA) using the binary matrix of present/absent gene-associated reactions across all 3070 models as a basis. X-axis: First principal component (PC1). Y-axis: Second principal component (PC2). % denotes the % of variance explained by the respective principal component. Strains are color-coded by their genetic lineage as described in Murray et al (24).

As our starting point, we examined the general properties of these models. We found that these models included ∼600 genes (corresponding to about one third of a typical *S. suis* genome, and in line with reconstructions from other bacterial species). Moreover, these models included ∼660 unique intracellular metabolites, and ∼830 gene-associated reactions, with limited strain-to-strain variability in the overall number of metabolites or reactions (**Figure 1B, Supplementary Figure 1**). Analysis of the pan-reactome identified ∼2000 unique gene-associated reactions across all 3070 models (**Figure 1C, Data set 2**), which showed a bimodal distribution: a smaller fraction of ∼670 reactions, corresponding to about 80% of reactions per model, were highly conserved (i.e. found in at least 90% of models), whereas the majority of reactions were only found in less than 10% of models (**Figure 1D, Supplementary Figure 2**). To test whether the reactions within these models are distributed in a lineage-dependent manner, we performed principal component analysis on the full pan-reactome matrix (that is, the matrix containing the gene-associated reactions which are present in each strain). As expected, given this high degree of reaction conservation, the first principal component captured most of the expected variance across models (∼90%, see **Figure 1E**). Nevertheless, models were still well separated according to different lineages when considering the first two principal components. This separation was particularly pronounced between the out-group lineages and the rest of the strains, but we also observed a separation of pathogenic and commensal lineages (including when removing the out-group strains, **Supplementary Figure 3**). Further analysis revealed a subset of 32 reactions that differed substantially in prevalence between pathogenic and commensal lineages (**Supplementary Figure 4**). Notably, among these reactions was a methylsulfonate transport reaction (BIGG reaction id: MSO3abc), which we had previously been linked to a pathogenic genomic island in *S. suis* (*24*). Thus, our findings suggested that at least some metabolic traits in *S. suis* are lineage dependent and vary across pathogenic versus commensal groups.

### Section 2: examining predicted amino acid requirements of S. suis

Next, we wanted to examine the predicted metabolic traits of these *S. suis* models in more detail. First, we focused on nutrient requirements, and specifically amino acid auxotrophies. An experimental study recently identified several amino acids for which a serotype 2 *S. suis* strain was auxotrophic *in vitro*, namely L-arginine, L-histidine, L-cysteine, and L-tryptophan (*39*). To test to which extent these auxotrophies are conserved across the various *S. suis* lineages, we performed Flux Balance Analysis (FBA) (*40*) to predict the impact of omitting individual amino acids on growth in a rich *in vitro* cultivation medium containing glucose as the only carbon source (see **Data set 3** for the approximated medium composition we used). In general, we found that the previously reported (*39*) amino acid auxotrophies for L-arginine, L-histidine, L-cysteine, and L-tryptophan, were highly prevalent in the different *S. suis* lineages (**Figure 2A** and **Data set 4**), with sporadic additional auxotrophies in few strains. These findings suggested that the nutrient requirements of *S. suis* are largely conserved. Nevertheless, the out-group lineages 2 and 3 diverged substantially from these auxotrophy patterns: out-group lineage 2 lacked most conserved auxotrophies except for L-cysteine, and out-group lineage 3 predicted to be auxotrophic for L-methionine, but not for L-histidine or L-tryptophan.

**Figure 2.**
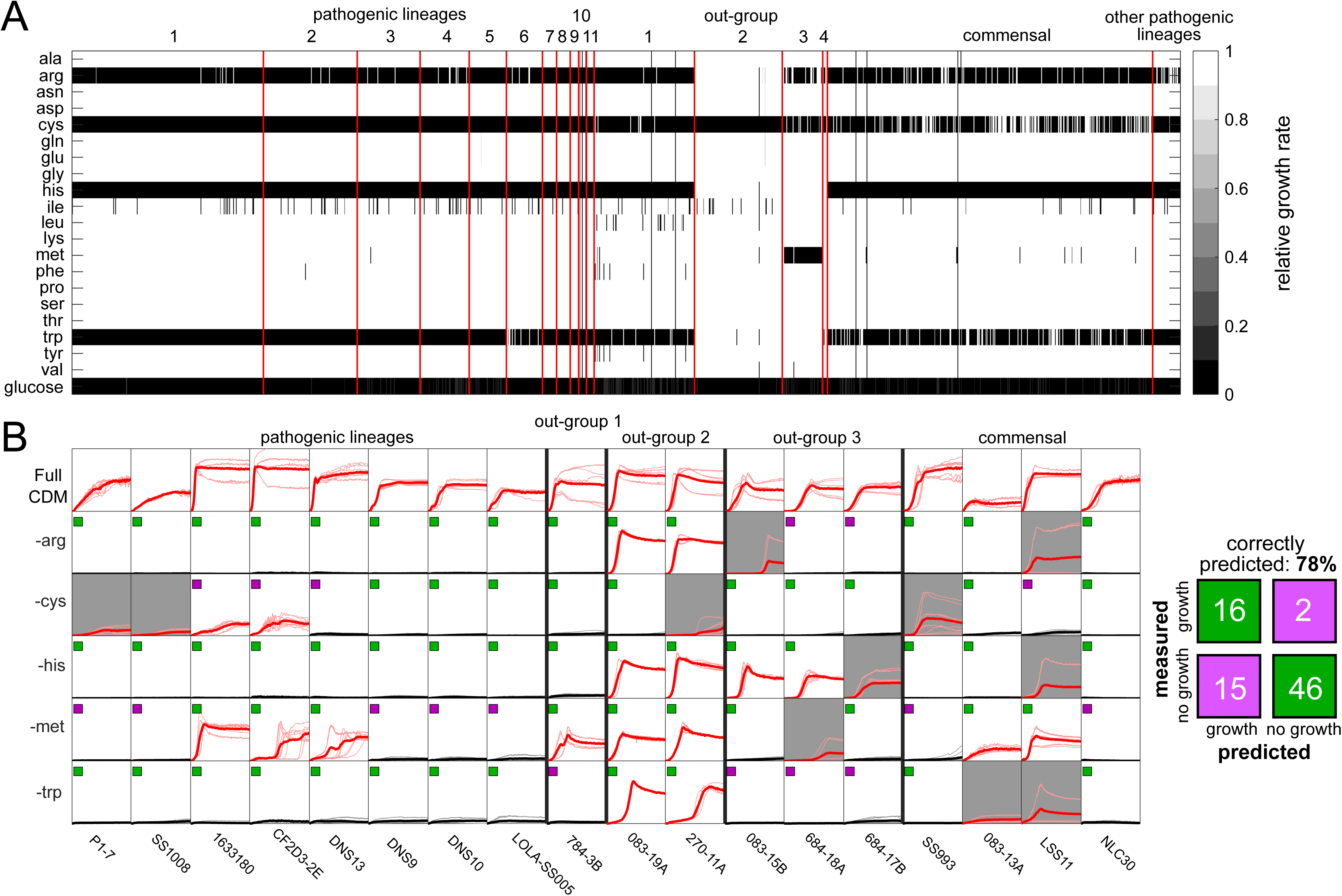
Predicting and experimentally validating nutrient auxotrophies in *S. suis* GSMM atlas. **A)** Simulated relative growth rate (compared to full medium) upon removing a single metabolite across all models. Metabolites were removed by setting the respective maximal uptake rate to 0. Each column denotes a strain-specific model, and models were sorted according to their lineage as described in Murray et al (24). **B)** Experimental validation using *in vitro* growth assays in chemically defined medium (CDM) for strains from different lineages. Data shown are time courses from growth experiments performed in 96-well plate format. Top column: growth in full CDM. Bottom columns: growth in CDM variants lacking individual amino acids, as indicated. Red: strain grew in selected condition (max OD600 >= 0.1). Black: strain did not grow in selected condition (max OD600 < 0.1). Thin lines: individual replicate cultures. Thick line: mean across replicate cultures (from 1-3 independent experiments and 3 replicates per experiment). Gray background: inconsistent growth across replicates (i.e. CV of max OD600 across replicates > 0.5) and therefore not considered in subsequent analyses. Small insets in each plot: comparison of experimental data with FBA prediction (green = matches prediction, purple = does not match prediction). X-axis range: 0-48 h. Y-axis range: 0-1.5 OD600. Right: corresponding confusion matrix comparing FBA predictions (x-axis) and experimental data (y-axis).

To validate these diverging auxotrophies experimentally, we performed amino acid drop out experiments in a chemically defined *in vitro* cultivation medium (*39*) using a selection of 18 strains from different lineages (**Figure 2B**). We found that the metabolic models predicted most amino acid auxotrophies in these strains correctly (i.e. 78% of amino acid-strain pairs were correctly predicted), suggesting that our GSMM atlas largely recapitulates the amino acid requirements of different *S. suis* lineages.

### Section 3: examining predicted carbon source repertoire of S. suis

The results shown above revealed lineage-dependent differences in amino acid auxotrophies in our GSMM atlas. However, beyond the external supply of amino acids, *S. suis* also requires access to a carbon source for growth: since *S. suis* lacks a full TCA cycle (*41*), it relies on glycolysis for ATP production. To examine whether the repertoire of carbon sources usable for growth is similarly lineage-dependent in *S. suis*, we next tested the ability of each strain to grow on 94 different glycolytic carbon sources. These included various mono-/di-/oligosaccharides, as well as sugar derivates, for which at least one of the strains was predicted to carry a transport reaction (see **Data set 5**). To account for differences in the number of carbon atoms, the maximal uptake of each carbon source was set to 60 mmol C / gCDW / h (corresponding to 10 mmol glucose / gCDW /h, the maximal uptake rate used in the amino acid auxotrophy simulations).

We found that among the tested carbon sources, only a subset of 34 were able to support growth in more than 10% of the strains (**Figure 3A**). Apart from glucose, these included several monosaccharides (e.g. fructose, mannose, galactose), some di-/trisaccharides (e.g. maltose, trehalose) as well as amino sugars such as N-acetyl-glucose. Nevertheless, there were substantial lineage-dependent differences in the number of utilizable carbon sources across strains: pathogenic lineages tended to have a larger carbon source range than out-group or commensal lineages (**Figure 3B**). A similar pattern emerged when categorizing strains not by lineage, but by disease association, with strains associated to systemic disease having a larger carbon source range (**Supplementary Figure 5**). These patterns were only weakly correlated with the number of genes or reactions present in each model (**Supplementary Figure 6**).

**Figure 3.**
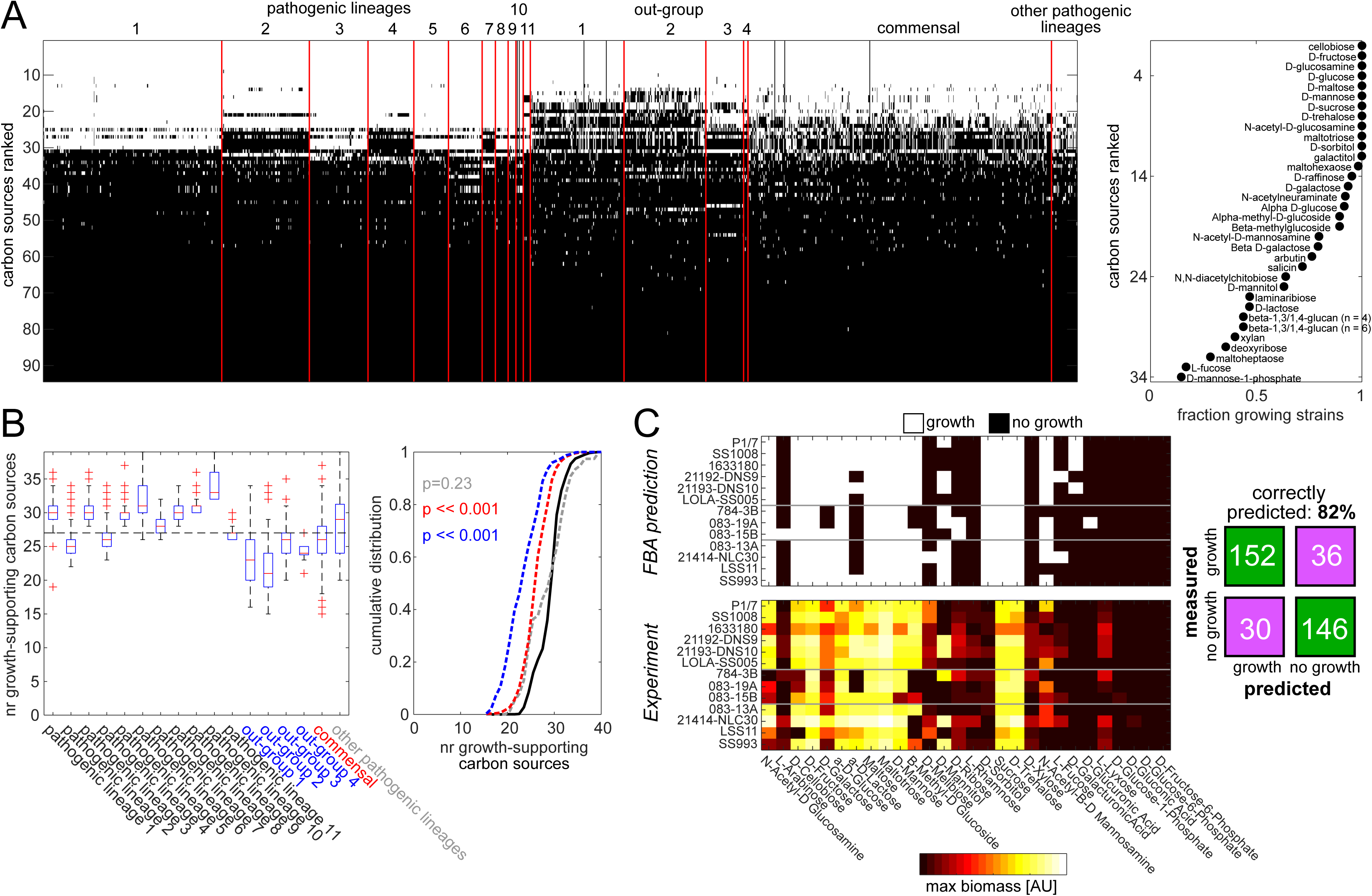
Carbon source utilization patterns in *S. suis* GSMM atlas. **A)** Left: Prediction of growth in 3070 strains (x-axis) on 94 single carbon sources (y-axis), ranked from top to bottom based on the fraction of strains that are predicted to grow in each carbon source (defined here as: growth rate reaches at least 10% of growth rate on glucose, see methods), and sorted by lineages. White: strain is predicted to grow in the respective carbon source. Right: same data but showing the fraction of strains predicted to grow on 34 carbon sources that are predicted to support growth in >10% of strains. **B)** Boxplots of the number of carbon sources predicted to support growth in each strain, split by lineages. Horizontal black line: median across all strains. Right, same data but shown as cumulative distributions. Black continuous line: Merged pathogenic lineages 1-11. Gray dashed line: other pathogenic lineages. Blue dashed line: merged out-groups 1-4. Red dashed line: commensal lineages. P-values show the result of a two-sided Wilcoxon rank sum test (testing whether the two sample distributions stem from continuous distributions with equal medians) in reference to the distribution in pathogenic lineages 1-11. **C)** Experimental validation of carbon source utilization in 13 strains (grouped into pathogenic lineages, out-groups, and commensals from top to bottom) and 28 carbon sources. Top: FBA-prediction. Bottom: maximal biomass production in Biolog plates (mean of three replicate plates, see methods). The corresponding growth curves are shown in **Supplementary** Figure 8. Right: corresponding confusion matrix comparing FBA predictions (x-axis) and experimental data (y-axis).

To experimentally validate these carbon utilization patterns, we used Biolog plates (i.e. PM1) and determined the ability of 13 strains from different lineages to metabolize 95 chemically diverse carbon sources. In line with our FBA predictions, only a small subset of these carbon sources (mostly sugars and sugar derivates) could be metabolized by any of the strains (**Supplementary Figure 7**). Importantly, when examining the 28 carbon sources for which we had both FBA predictions and experimental data, the match between predictions and experimental results was high (i.e. correct prediction in 82% of the cases, **Figure 3C, Supplementary Figure 8**). While for most of these carbon sources strains from different lineages tended to behave similarly in both FBA predictions and experiments, there were exceptions. In particular, our experimental data suggested that beta-methyl-glucoside preferentially supported the growth of strains from pathogenic lineages, although that difference was less pronounced in the FBA predictions (**Figure 3C**). Using a larger panel of 72 *S. suis* strains, we confirmed that the ability to utilize beta-methyl-glucoside as a carbon source in chemically defined media is more prevalent in pathogenic lineages than out-group/commensal strains (**Supplementary Figure 9**). Thus, these experimental data support the notion that, although carbon utilization patterns are largely conserved across strains, pathogenic lineages tend to have a broader carbon source range.

### Section 4: Predicted S. suis growth and reaction essentiality in inferred in vivo environments

The data described above suggested that while metabolic traits are largely conserved in *S. suis*, there are nevertheless some lineage-dependent differences. Next, we wanted to examine the potential *in vivo* impact of these differences. Towards this end, we decided to assess the simulated ability of each strain to grow in different plausible *in vivo* niches, namely the nasal environment (i.e. the normal habitat of *S. suis*), as well as plasma and cerebrospinal fluid (CSF) as environments where *S. suis* can be found during systemic infections (*5*, *42*). Since there was little information available on the metabolites present in these environments in pigs, we used *in vivo* data from humans to approximate their composition (*43*, *44*) (see **Figure 4A** and **Data set 2** for the approximated composition of each environment). We found that the vast majority of models predicted growth in all tested *in vivo* environments, regardless of lineage or disease association (**Figure 4B, Supplementary Figure 10**). An exception was out-group lineage 3, which was unable to grow in those environments that lack L-methionine (presumably due to its additional L-methionine auxotrophy, see section 2).

**Figure 4.**
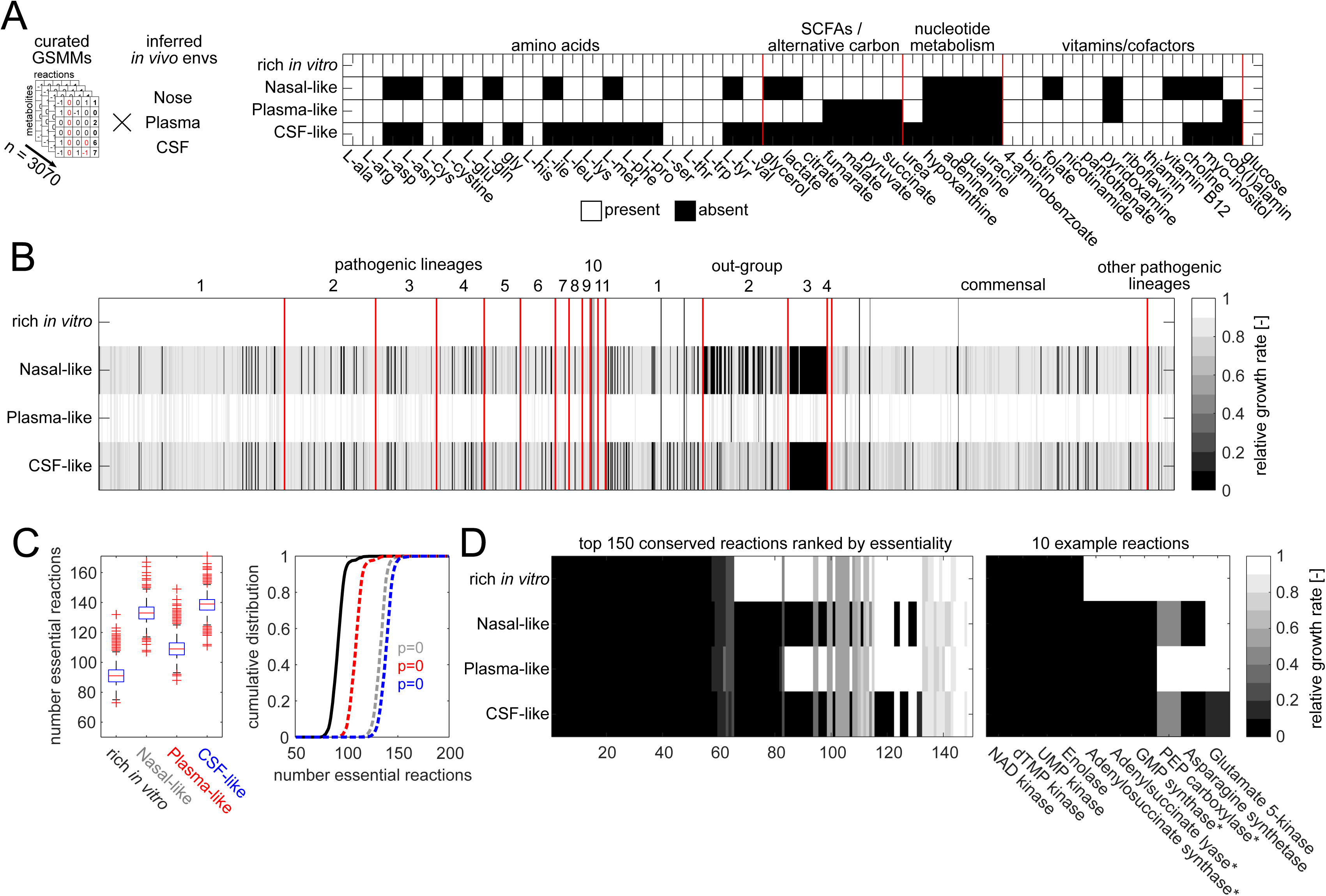
Predicted growth of *S. suis* GSMM atlas in inferred *in vivo* environments. **A)** Schematic of approach, and composition of simulated *in vivo* environments (inferred from *in vivo* measurements in humans, see main text). **B)** Predicted growth (relative to a rich *in vitro* condition) in each simulated *in vivo* condition across all strains. **C)** Left: Boxplots of the number of essential reactions (i.e. reactions whose deletion leads to a relative growth rate below 0.001) across all models in each tested condition. Right: same data but shown as cumulative distributions. Black continuous line: rich *in vitro* condition. Gray dashed line: Nasal-like condition. Red dashed line: Plasma-like condition. Blue dashed line: CSF (cerebrospinal fluid). P-values show the result of a two-sided Wilcoxon rank sum test. **D)** Left: Mean growth rate (across all strains) in the 150 conserved reactions (i.e. reactions present in >90% of strains) with the largest overall effect on essentiality. Right: 10 example reactions. * denote reactions associated with genes that were shown to be conditionally essential during *in vivo* pig infections (9).

Although most strains in our GSMM atlas were predicted to grow in all tested *in vivo* environments, we hypothesized that these environments (which contain fewer metabolites than the simulated rich *in vitro* condition, see **Figure 4A**) may nevertheless cause some metabolic reactions to become conditionally essential (that is, essential *in vivo* but not *in vitro*). To test this conjecture, we systematically predicted the impact of deleting individual reactions in each model-condition pair (in total, 15+ million unique model-reaction-condition combinations). As expected, the inferred nasal and CSF environments, which have the fewest metabolites, showed significantly higher numbers of essential reactions (i.e. reactions whose deletion reduces the relative growth rate below 0.001) across the tested models (**Figure 4C**).

To examine these essential reactions in more detail, we focused on 677 highly conserved reactions (i.e. found in > 90% of strains) and calculated the mean growth rate upon their deletion in each condition (**Figure 4D** and full list in **Data set 6**). A subset of 65 reactions was predicted to be essential in all four tested conditions (i.e. *in vivo* and *in vitro*). Reassuringly, these included many reactions known to be essential in streptococci and other bacterial species, such as NAD kinase, dTMP kinase (*45*), UMP kinase (*46*), and reactions of lower glycolysis (e.g. enolase, phosphoglycerate kinase). Another 17 reactions were predicted to be essential in all three inferred *in vivo* environments, but not *in vitro* (**Supplementary Table 3**). These were largely reactions belonging to purine metabolism, such as adenylosuccinate synthase (**PurA**), adenylsuccinate lyase (**PurB**), and GMP synthase (**GuaA**), all three of which were recently shown experimentally to be conditionally essential during *in vivo* pig infections (*9*). Another subset of ∼50 reactions was essential only in two or one of the tested *in vivo* conditions. For example, glutamate 5-kinase, which is part of the glutamate biosynthesis pathway, was only essential in CSF, where glutamate is predicted to be absent. Overall, these findings suggest the existence of a subset of metabolic reactions in *S. suis* that are dispensable during growth in rich *in vitro* conditions but become conditionally essential *in vivo*.

## Discussion

In this study, we generated the first genome-scale metabolic model atlas of the zoonotic pathogen *Streptococcus suis*, with 3070 strain-specific metabolic models that cover the whole breadth of currently known lineages. Using this GSMM atlas, we examined three key metabolic traits of *S. suis* at large-scale, namely amino acid auxotrophy, the ability to use different carbon sources, and the ability to grow in plausible *in vivo* environments.

We found that in general, the tested metabolic traits within this GSMM atlas are largely similar across *S. suis* strains and lineages. This conservation not only manifests in terms of amino acid auxotrophy (**Figure 2**), but also carbon source range (**Figure 3**). This finding is in line with genomic analyses from other nasal colonizers (*47*), and suggests that despite the substantial genetic variability found in *S. suis*, its **metabolism** is nevertheless largely conserved at the functional level. An important consequence of this high degree of metabolic conservation is that both pathogenic and non-pathogenic lineages are in principle equipped to grow in different simulated *in vivo* environments (**Figure 4**), suggesting that *in vivo* metabolite availability *per se* is a weak barrier against systemic infection in most *S. suis* lineages.

Nevertheless, we did detect important differences in metabolic traits in at least some of the lineages. The clearest example are divergent amino acid auxotrophies in the out-group strains (**Figure 2**), which suggest that these strains may in fact occupy a different environmental niche *in vivo*. Currently we do not know what this niche may be. One possibility is that these out group strains are naturally colonizing the porcine vagina instead, since genomic comparison of newly isolated porcine vaginal *Streptococcus* species revealed that they are closely related to *S. suis* (*48*). In line with this hypothesis, recent genomic studies have identified distinct subgroups among isolates that all had previously been classified as *S. suis*, suggesting the existence of a broader *Streptococcus suis* complex (*49*, *50*) that may include lineages with distinct niches. Future efforts may use our findings as a starting point for elucidating the relationship between metabolic traits and *in vivo* niches among the members of this *Streptococcus suis* complex.

Another lineage-dependent difference we detected was the larger range of growth-supporting carbon sources in pathogenic *S. suis* lineages (**Figure 3**), in line with previous genomic analyses (*23*). One caveat is that accurately matching transporters to carbon sources from genomic information (which is also at the heart of the GSMM reconstruction method we used) remains highly challenging. Therefore, it is conceivable that the annotation of carbon source transporters in our GSMM atlas is in some cases incorrect. For example, we predict that pathogenic lineages can use chitobiose (a chitin building block) as a carbon source, which is unlikely to be present at large quantities in porcine or human hosts (but is nevertheless structurally related to mucin building blocks). Another example is beta-methyl-glucoside: while the FBA predictions suggested very few differences in the ability to utilize this sugar across strains (**Figure 3**), our experimental data does indicate that in fact beta-methyl-glucoside utilization is much more prevalent in pathogenic lineages (**Supplementary Figure 9**), making it an attractive phenotypic marker e.g. for diagnostic purposes. Future efforts may use our findings to further refine and experimentally validate the carbon source range of *S. suis* lineages.

Finally, by systematically predicting the essentiality of reactions within our GSMM atlas in different *in vitro* and *in vivo* conditions, we were able to shed light on the role of potential metabolic virulence factors (**Figure 4**). Specifically, our analyses identified several reactions in purine metabolism, including adenylosuccinate synthase (**PurA**), adenylsuccinate lyase (**PurB**), and GMP synthase (**GuaA**), as *in vivo* metabolic vulnerabilities. Interestingly, an *in vivo* experiment using transposon libraries recently demonstrated that deletion mutants of the associated genes have massively impaired fitness during pig infection (*9*), and PurA was identified to be conditionally essential during growth in porcine serum (*51*) in line with our predictions. Thus, it is tempting to speculate that at least a portion of the metabolic genes that were previously identified as virulence factors (often using gene deletion experiments) are not direct drivers of pathogenicity. While they fit a broad definition of enabling *in vivo* survival, they act as conditionally essential fitness factors sustaining basic *in vivo* metabolic needs rather than actively driving disease. Interestingly, purine biosynthesis has been identified as a metabolic vulnerability in various other pathogens (*52*). Future efforts may leverage our GSMM atlas to guide the design of new antimicrobials that target purine biosynthesis (e.g. through specific enzyme inhibitors) in *S. suis*.

One caveat is that in some cases our predictions did not match experimental evidence. For example, the aforementioned *in vivo* transposon library experiment (*9*) also identified several reactions in carbon metabolism to be conditionally essential *in vivo*, most of which (with the exception of PEP carboxylase, see **Figure 4**) had no impact on *in vivo* growth in our simulations. Future efforts may explore these discrepancies to further refine computational predictions made by our GSMM atlas.

This study has several limitations. First, to enable the reconstruction and characterization of 3000+ metabolic models, we used an automated model reconstruction and curation approach. Although our experimental validation data do suggest that these models recapitulate at least some important metabolic traits, it is likely that our automated model reconstruction approach does not take all strain differences into account. For example, we did not account for potential differences in the composition of the cell capsule, which can differ substantially across *S. suis* serotypes (*53*). Future studies may further refine the models developed in this GSMM atlas to incorporate additional information if needed.

Second, to predict the ability of the strains in our GSMM atlas to grow *in vivo*, we approximated different plausible *in vivo* environments (i.e. nose, plasma, CSF) based on available information from humans (*43*, *44*). It is well possible that the exact metabolites available in these environments may be different in the porcine host, but currently such *in vivo* data are scarce in pigs. Nevertheless, we note that some of the metabolic reaction we predict to be conditionally essential *in vivo*, in particular reactions belonging to purine metabolism (see **Figure 4**), have shown to be also conditionally essential in *S. suis* in *in vivo* (*9*, *54*) or *in vitro* (*51*) experiments, suggesting that our simulations do recapitulate at least some physiologically relevant properties of the *in vivo* porcine metabolic environment. Future efforts may quantify the metabolic composition of these environments in pigs in more detail to refine the prediction of *in vivo* essential metabolic reactions.

Third, our predictions are limited to metabolic traits that manifest as differences in biomass production. Therefore, we will miss differences in metabolic traits that do not affect biomass production directly. For example, consistent with previous genomic analyses (*12*), we found that pathogenic *S. suis* lineages lack most reactions needed for the *de novo* biosynthesis of polyamines (see **Supplementary Figure 2**). However, while there is clear evidence that polyamines affect the *in vivo* virulence of *S. suis* (*9*) and other Streptococci (*55–58*), polyamines are not directly connected to biomass production. Therefore, even *S. suis* mutants completely devoid of polyamines can grow at least *in vitro* (*12*). As the exact role of polyamines in bacterial physiology remains enigmatic (*59*), future efforts will be needed to clarify the mechanisms by which polyamine metabolism affects *S. suis* virulence *in vivo*.

In conclusion, by mapping the metabolic landscape of the entire *S. suis* complex, we demonstrate that metabolism is not a limiting factor for invasive infection, rather, the pathogenicity of *S. suis* appears linked to increases in metabolic flexibility in carbon use, enabling the pathogen to better exploit the host environment. This work also has direct translational applications. We identify specific *in vivo* vulnerabilities, such as purine usage, as potential targets for vaccine design, and biochemical pathways over-represented in pathogenic lineages to increase diagnostic accuracy when screening healthy pigs. Ultimately, these findings provide a model for understanding how metabolic niche facilitates the transition from commensalism to pathogenicity, providing new tools to predict and control emerging pathogenic strains of *S. suis*.

## Methods

### Model reconstruction and curation

Genome assemblies and functional annotations for 3,070 *S. suis* strains were obtained from a previous study (*24*). In that study, all raw reads sequenced via Illumina technology were processed through a pipeline using SPAdes v.3.12.0 (*60*) for assembly and Prokka v.1.14.5 (*61*) for annotation. From each strain, a genome-scale metabolic model was reconstructed using CarveMe (*36*) (Version 1.6.1 downloaded from https://github.com/cdanielmachado/carveme in April of 2024). CarveMe was used as provided, with one small modification: to ensure that as many metabolic genes as possible were considered during the reconstruction, the annotation process was slightly relaxed by removing a hard-coded cut-off value for the blast annotation score (originally set to 100 in this version of CarveMe). Each model was reconstructed using the following parameters: First, since currently there is no experimental data available on the exact biomass composition of *Streptococcus suis*, the biomass composition in each model was approximated using a universal gram-positive biomass vector (*36*). Second, a gap filling step was included using the composition of a plausible rich cultivation medium, which closely matches the composition of a chemically defined medium that supports *S. suis* growth *in vitro* (*39*) (see **Data set 2**). MEMOTE (*38*) was then used to assess the overall quality of each reconstructed model (we obtained MEMOTE scores between 70 and 90% for all models, see **Data set 1**). All these steps were performed on a standard laptop (64-bit Linux system, with an Intel Core i7-10750H CPU and 32 GB of RAM).

Each model was further curated based on experimental evidence. First, based on ^13^C-labeling data from a recent *in vitro* study (*39*), the directionality of glutamate dehydrogenase (BIGG reaction ID GLUDy) was restricted, only allowing the consumption, but not the production of glutamate (by setting its upper bound to 0). Second, based on *in vitro* exometabolomics data, which demonstrated that *S. suis* can consume adenine in rich *in vitro* cultivation media (*62*), an adenine transport reaction was included in those models that did not already have it. Third, to account for the fermentation products previously reported for *S. suis* (i.e. lactate, acetate, ethanol, see (*41*)) transport reactions for these metabolites were included in all models that did not already have them.

In addition, further curation steps were included based on initial simulation results. First, several redundant reactions around the pyruvate node were removed (BIGG reaction IDs PEPC, PYK6, PDHa, PFL). Second, in initial simulations substantial secretion of several unexpected and highly unlikely metabolites (i.e. BIGG metabolite IDs 12ppd R, 2h3mb, 2h3mp, 2mpa, 3mba, 2mba) was observed. To avoid this simulation artifact, the secretion of these metabolites was blocked, and the directionality of another reaction was restricted (BIGG reaction ID PAPSSH, upper bound set to zero). All these curation steps were implemented in an automated manner using custom MATLAB scripts and commands from the Cobra toolbox (*63*) (Version 3.4, downloaded from https://github.com/opencobra/cobratoolbox in April of 2024).

### Model simulations

All flux balance analysis (FBA) simulations were performed using custom MATLAB scripts and commands from the Cobra toolbox (*63*) (Version 3.4, downloaded from https://github.com/opencobra/cobratoolbox in April of 2024), and using Gurobi (Version 9.1.2) as the default solver. Briefly, nutrient auxotrophy simulations were performed by setting the maximal uptake rate of the respective metabolite to zero and predicting the maximal growth rate using the Cobra toolbox command optimizeCbModel(model, ‘max’). Similarly, carbon source utilization simulations were performed by setting the maximal uptake rate of the respective carbon source 60 mmol Carbon / gCDW /h to account for different numbers of carbon atoms in these carbon sources, and predicting the maximal growth rate using the Cobra toolbox command optimizeCbModel(model, ‘max’). Only cases in which a model has the respective transport reaction were considered (the rest were set to the maximal growth rate in absence of any carbon sources). Relative growth rates for each carbon source-strain pair (compared to growth on glucose) were calculated as μ_relative_ = (μ_carbon source_ – μ_no carbon_)/ (μ_glucose_ – μ_no carbon_) to account for the fact that in some cases the predicted maximal growth rate without a carbon source was not exactly zero. If μ_relative_ was above 0.001, the respective carbon source-strain pair was defined as “growing”. Finally, reaction essentiality in each inferred *in vivo* environment was determined by first setting the boundaries of the included exchange reactions as listed in **Data set 2**, and then using the Cobra toolbox to calculate the relative growth rate of each predicted reaction deletion (command singleRxnDeletion(model)).

All simulation results were processed and visualized using custom MATLAB scripts (Version R2021a). All code needed to recreate the analyses and plots, as well as the resulting data sets, are provided below under the section **Data availability**.

### Experimental validation by growth assays in chemically defined medium

Experimental validation of predicted metabolic traits was performed using amino acid and sugar auxotrophy assays in which *S. suis* strains were grown in chemically defined medium (CDM) with selective omission or replacement of target amino acids or carbohydrates as described below.

CDM was prepared as previously described by Willenborg et al (*39*) with minor modifications. The CDM consisted of a basal medium, a defined amino acid mixture, and glucose as the primary carbon source, unless stated otherwise. The basal medium contained salts, vitamins, nucleotides, and cofactors. First, Na₂HPO₄·2H₂O, folic acid, riboflavin, adenine, guanine, and uracil were dissolved together in 600 mL sterile deionized water. All remaining components were prepared separately in sterile deionized water and added at the required volumes. The final volume was adjusted to 900 mL with sterile deionized water, followed by filter sterilization through a 0.22 µm membrane unit (Steritop, Millipore). Prepared basal medium was stored at 4 °C until use. Stock solutions of 21 amino acids were prepared at the required concentrations using sterile water, 1 M HCl, or 1 M NaOH, depending on solubility. Glucose or other carbohydrates were prepared separately as a 1.5 M stock solution in sterile deionized water. All stock solutions were filter sterilized and stored at 4 °C. The exact CDM composition used here, and the concentrations of each stock solution are listed in **Supplementary Table 1**.

For amino acid auxotrophy testing, 18 strains were selected from different lineages. The six amino acids tested were arginine, cysteine/cystine, histidine, methionine, and tryptophan. To test the ability of *S. suis* strains to utilize beta-methyl-glucoside, 72 strains were selected from different lineages, and glucose was included as the default carbon source and positive growth control. The exact strains used in these experiments are listed in **Supplementary Table 2**.

Growth assays using these strains in CDM were performed as follows. Bacterial strains were streaked onto Columbia Blood Agar plates (Oxoid) and incubated overnight at 37 °C under aerobic conditions. The following day, overnight broth cultures were initiated from one to three single colonies in Todd Hewitt broth (THB) supplemented with 0.2% yeast extract (Oxoid). Cultures were incubated overnight and then diluted to an optical density at 600 nm (OD_600_) of 0.02 in 10 ml fresh THB containing 0.2% yeast extract. Cells were grown for approximately 4 to 5 hours to mid-exponential phase. For inoculation into CDM, 10 mL of each culture was pelleted (10 min ,4,000 rpm, 4°C) and washed with 4 mL CDM basal medium to remove residual rich medium. Cell pellets were then resuspended in 5 mL CDM basal medium and then adjusted to an OD_600_ of 0.02 in either complete CDM or modified CDM formulations lacking selected amino acids or containing alternative carbohydrates. Subsequently, aliquots of 200 µL were dispensed into 96-well microplates together with medium-only controls. Growth was monitored in duplicate or triplicate using an automated plate reader (BIOSCREEN, Finland) with OD_600_ measurements recorded every 30 min for 48 h (amino acid experiments) or 24 h (sugar utilization experiments) at 37 °C with continuous moderate shaking.

Data analysis was performed in R (version 4.4.2) and using custom MATLAB scripts (Version R2021a).

### Experimental validation by Biolog growth assays

In addition, carbon utilization patterns of 13 *S. suis* strains from different lineages were determined using Biolog plates (i.e. PM1 plates with 95 different carbon sources, Catalog #12111). All biolog reagents were supplied by TECHNOPATH Distribution Ltd. In brief, bacterial strains were streaked onto Columbia Blood Agar plates (Oxoid) and incubated overnight at 30 °C under aerobic conditions. The following day, cells were scrapped using a loop and added to inoculation fluid IF-0a (Catalog# 72268) to an OD600 (600 nm) of 0.04, to create the initial cell suspension. Experimental inoculation fluid was set up following manufacturer’s instructions (00A 121 Rev A Gram Positive Phenotype Microarray Protocol and 00A 129 Rev A - Phenotypic MicroArray Alternate Protocol Appendix with minor alterations detailed below). Per sample 12 mL of final inoculation fluid was prepared containing, 10 ml of 1.2x IF-0a, 1 mL of PM additive solution (PM additive solution - (per 100 mL, 10 mL (240 mM MgCl2 6H2O and 120 mM CaCl2 2H2O), 10 mL (0.6 mM L-glutamate, Na) 30 mL (0.5 mM L-cystine pH 8.5 and 1 mM 5’-UMP, 2Na), 10 mL (3mM, L-Arginine HCl, 3 mM Hypoxanthine, 0.6 mM β-NAD hydrate, 30 µM Riboflavin, 0.6% Yeast extract, 0.6% Tween 80, 240 mM Na2HPO4 pH 6.0, L-methionine 3 mM and 1.2 mM alpha-lipoic acid) and 40 mL of sterile dH2O – PM additive solution end), 0.88 mL of initial cell suspension and 0.12 mL of dye mix G (Catalog# 74227). To each PM1 plate well, 100 µL of final inoculation fluid was added. Biomass production was monitored at 30°C over 66h using an OmniLog automated plate reader and incubator. Each strain was tested in triplicate. Data were extracted using the Biolog software (conversion of D5E to OKA: D5E_OKA Data File Converter v1.1.1.15 and extraction of raw kinetic data using PM analysis software: Kinetic V1.3). Subsequently, growth curves were analyzed in custom MATLAB scripts (R2021a) as follows: Data were normalized by subtracting the signal at the first time point, and the maximal biomass signal for each replicate was identified within the 66h cultivation curve. Subsequently, background signals were removed by subtracting the mean maximal biomass signal of the negative control (i.e. no carbon source).

## Data availability

Models and the MATLAB code needed to reproduce analyses and figures are available in the CORA repository (https://doi.org/10.34810/DATA3435).

Data sets available with this study:

- **Data set 1**: Metadata for all models, accompanying model features (i.e. number of genes, metabolites, gene-associated reactions, reactions, gapfilled reactions), and MEMOTE scores
- **Data set 2**: Composition and exchange boundaries for simulated conditions used here
- **Data set 3**: List of reactions in pan-reactome and their associated genes in each strain
- **Data set 4**: Predicted relative growth rate of each strain in auxotrophy simulations
- **Data set 5**: Predicted relative growth rate of each strain in carbon source utilization simulations
- **Data set 6**: Predicted mean relative growth rate when deleting (one at a time) 677 conserved reactions in all tested conditions

## Supplementary information

Supplementary information is provided as a single document containing:

- Supplementary tables 1-3
- Supplementary figures 1-10

## Funding

This work was supported by funding from the Spanish Ministry of Research and Innovation (RYC2021-033035-I/AEI/10.13039/501100011033 and PID2023-152210NA-I00//AEI/10.13039/501100011033 to KK). PO was supported by an FPU (FPU19/02126) fellowship. APF was funded by a fellowship from the European Union’s Horizon 2020 Research and Innovation Program under the Marie Sklodowska-Curie Grant Agreement number 956154, as part of the Innovative Training Network (Innovative Approaches to Identification of Metabolic Targets for Antimicrobials [INNOTARGETS]) (https://cordis.europa.eu/project/id/956154). PAH acknowledges funding from BBSRC (BB/T001038/1, BB/Y00082X/1) and the Royal Academy of Engineering Research Chair Scheme for long term personal research support (RCSRF2021\11\15). This work was also supported by Innovate UK project 10101990.

## Author contributions

Conceived and designed the study: KK, LW. Performed experiments: CL, MD, IL, HW, JTM, APF. Performed analyses: KK, CL, PO, GM, JTM, DM, LW. Supervised experiments and analyses: KK, PAH, FCF, VA, AWT, LW. Wrote manuscript with contributions from all authors: KK.

## Conflict of Interest

The authors declare no conflict of interest.

## Supporting information

Data set 1

Data set 2

Data set 3

Data set 4

Data set 5

Data set 6

## Key abbreviations

GSMM: genome-scale metabolic model
FBA: flux balance analysis
CDM: chemically defined medium
CSF: cerebrospinal fluid

## Notes

https://doi.org/10.34810/DATA3435

### Competing Interest Statement

The authors have declared no competing interest.

