## Supplementary Information for "Genome-scale metabolic model atlas of the zoonotic pathogen *Streptococcus suis*"

### Supplementary Tables

**Supplementary Table 1. Strains used in the different experimental validation assays**

| **Strain ID** | **Lineage/Classification** | **Validation assay** |
| --- | --- | --- |
| P1_7 | Pathogenic lineage 1 | Amino acid auxotrophy |
| SS1008 | Pathogenic lineage 1 | Amino acid auxotrophy |
| 1633180 | Pathogenic lineage 3 | Amino acid auxotrophy |
| CF2D3-2E | Pathogenic lineage 3 | Amino acid auxotrophy |
| 21196_DNS13 | Pathogenic lineage 4 | Amino acid auxotrophy |
| 21192_DNS9 | Pathogenic lineage 5 | Amino acid auxotrophy |
| 21193_DNS10 | Pathogenic lineage 5 | Amino acid auxotrophy |
| LOLA_SS005 | Pathogenic lineage 6 | Amino acid auxotrophy |
| 784_3B | out-group 1 | Amino acid auxotrophy |
| 083_19A | out-group 2 | Amino acid auxotrophy |
| 270_11A | out-group 2 | Amino acid auxotrophy |
| 083_15B | out-group 3 | Amino acid auxotrophy |
| 684_18A | out-group 3 | Amino acid auxotrophy |
| 684_17B | out-group 3 | Amino acid auxotrophy |
| SS993 | Commensal | Amino acid auxotrophy |
| 083_13A | Commensal | Amino acid auxotrophy |
| LSS11 | Commensal | Amino acid auxotrophy |
| 21414_NLC30 | Commensal | Amino acid auxotrophy |
| P1_7 | Pathogenic lineage 1 | Carbon source (Biolog) |
| SS1008 | Pathogenic lineage 1 | Carbon source (Biolog) |
| 1633180 | Pathogenic lineage 3 | Carbon source (Biolog) |
| 21192_DNS9 | Pathogenic lineage 5 | Carbon source (Biolog) |
| 21193_DNS10 | Pathogenic lineage 5 | Carbon source (Biolog) |
| LOLA_SS005 | Pathogenic lineage 6 | Carbon source (Biolog) |
| 784_3B | out-group 1 | Carbon source (Biolog) |
| 083_19A | out-group 2 | Carbon source (Biolog) |
| 083_15B | out-group 3 | Carbon source (Biolog) |
| 083_13A | Commensal | Carbon source (Biolog) |
| 21414_NLC30 | Commensal | Carbon source (Biolog) |
| LSS11 | Commensal | Carbon source (Biolog) |
| SS993 | Commensal | Carbon source (Biolog) |
| 1148794 | Pathogenic lineages | Beta-methyl-glucoside test |
| 1508445 | Pathogenic lineages | Beta-methyl-glucoside test |
| 1522228 | Pathogenic lineages | Beta-methyl-glucoside test |
| 1602949 | Pathogenic lineages | Beta-methyl-glucoside test |
| 1619952 | Pathogenic lineages | Beta-methyl-glucoside test |
| 1633180 | Pathogenic lineages | Beta-methyl-glucoside test |
| 1637946 | Pathogenic lineages | Beta-methyl-glucoside test |
| M101830_S3 | Pathogenic lineages | Beta-methyl-glucoside test |
| 19787_M102942_S9 | Pathogenic lineages | Beta-methyl-glucoside test |
| M102942_S11 | Pathogenic lineages | Beta-methyl-glucoside test |
| M107642_R50 | Pathogenic lineages | Beta-methyl-glucoside test |
| 684_6A | Pathogenic lineages | Beta-methyl-glucoside test |
| DB1V3_4A | Pathogenic lineages | Beta-methyl-glucoside test |
| DNS1 | Pathogenic lineages | Beta-methyl-glucoside test |
| DNS10 | Pathogenic lineages | Beta-methyl-glucoside test |
| DNS2 | Pathogenic lineages | Beta-methyl-glucoside test |
| DNS31 | Pathogenic lineages | Beta-methyl-glucoside test |
| DNS9 | Pathogenic lineages | Beta-methyl-glucoside test |
| LOLA_SS009 | Pathogenic lineages | Beta-methyl-glucoside test |
| LSS70 | Pathogenic lineages | Beta-methyl-glucoside test |
| LSS73 | Pathogenic lineages | Beta-methyl-glucoside test |
| MY1C3_3B | Pathogenic lineages | Beta-methyl-glucoside test |
| NLR11 | Pathogenic lineages | Beta-methyl-glucoside test |
| NLR35 | Pathogenic lineages | Beta-methyl-glucoside test |
| NLS26 | Pathogenic lineages | Beta-methyl-glucoside test |
| NLS4 | Pathogenic lineages | Beta-methyl-glucoside test |
| NLS47 | Pathogenic lineages | Beta-methyl-glucoside test |
| NLS50 | Pathogenic lineages | Beta-methyl-glucoside test |
| P1_7 | Pathogenic lineages | Beta-methyl-glucoside test |
| PH1993199812 | Pathogenic lineages | Beta-methyl-glucoside test |
| PH1993199822 | Pathogenic lineages | Beta-methyl-glucoside test |
| PH1993199867 | Pathogenic lineages | Beta-methyl-glucoside test |
| SS1008 | Pathogenic lineages | Beta-methyl-glucoside test |
| Y02125 | Pathogenic lineages | Beta-methyl-glucoside test |
| 083_13A | Other lineages | Beta-methyl-glucoside test |
| 083_15B | Other lineages | Beta-methyl-glucoside test |
| 083_19A | Other lineages | Beta-methyl-glucoside test |
| 083_22A | Other lineages | Beta-methyl-glucoside test |
| 083_1C | Other lineages | Beta-methyl-glucoside test |
| 1129711 | Other lineages | Beta-methyl-glucoside test |
| 1191316 | Other lineages | Beta-methyl-glucoside test |
| 1197165 | Other lineages | Beta-methyl-glucoside test |
| 1466571 | Other lineages | Beta-methyl-glucoside test |
| 1628469 | Other lineages | Beta-methyl-glucoside test |
| 19813_M106115_S35 | Other lineages | Beta-methyl-glucoside test |
| 19843_M103975_R15 | Other lineages | Beta-methyl-glucoside test |
| M107545_R48 | Other lineages | Beta-methyl-glucoside test |
| 784_3B | Other lineages | Beta-methyl-glucoside test |
| DNC16 | Other lineages | Beta-methyl-glucoside test |
| DNC22 | Other lineages | Beta-methyl-glucoside test |
| DNC27 | Other lineages | Beta-methyl-glucoside test |
| DNC8 | Other lineages | Beta-methyl-glucoside test |
| DNR11 | Other lineages | Beta-methyl-glucoside test |
| DNR18 | Other lineages | Beta-methyl-glucoside test |
| DNR29 | Other lineages | Beta-methyl-glucoside test |
| DNR31 | Other lineages | Beta-methyl-glucoside test |
| DNS50 | Other lineages | Beta-methyl-glucoside test |
| LSS11 | Other lineages | Beta-methyl-glucoside test |
| LSS16 | Other lineages | Beta-methyl-glucoside test |
| LSS36 | Other lineages | Beta-methyl-glucoside test |
| LSS53 | Other lineages | Beta-methyl-glucoside test |
| LSS78 | Other lineages | Beta-methyl-glucoside test |
| LSS79 | Other lineages | Beta-methyl-glucoside test |
| LSS91 | Other lineages | Beta-methyl-glucoside test |
| NLC30 | Other lineages | Beta-methyl-glucoside test |
| NLR32 | Other lineages | Beta-methyl-glucoside test |
| NLR49 | Other lineages | Beta-methyl-glucoside test |
| NLR5 | Other lineages | Beta-methyl-glucoside test |
| NLR8 | Other lineages | Beta-methyl-glucoside test |
| PH1993199814 | Other lineages | Beta-methyl-glucoside test |
| SS993 | Other lineages | Beta-methyl-glucoside test |
| TMW_SS028 | Other lineages | Beta-methyl-glucoside test |

**Supplementary Table 2. Composition of chemically defined medium (CDM). Black**: media components that are weighed in together and used to prepare the base medium. **Green**: media components that are prepared in separate stock solutions. **Red**: amino acids that are prepared in separate stock solutions. **Purple**: carbon source (glucose for standard CDM).

| **Compound** | **Final concentration in CDM [mg/L]** | **Stock concentration and solvent**  **(if applicable)** |
| --- | --- | --- |
| FeSO_4_ x 7H_2_O | 10 | 1000x (in H2O) |
| Fe(NO_3_)_2_ x 9H_2_O | 2 | 1000x (in H2O) |
| K_2_HPO_4_ | 400 | 100x (in H2O) |
| KH_2_PO_4_ | 2,000 | 100x (in H2O) |
| MgSO_4_ x 7H_2_O | 682 | 100x (in H2O) |
| MnSO_4_ | 10 | 1000x (in H2O) |
| L-Alanine | 100 | 1000x (in H2O) |
| L-Arginine | 100 | 1000x (in H2O) |
| L-Aspartic acid | 100 | 250x (in 1M NaOH) |
| L-Asparagine | 100 | 200x (in H2O) |
| L-Cysteine | 100 | 500x (in 1M HCl) |
| L-Cystine | 100 | 500x (in 1M HCl) |
| L-Glutamic acid | 100 | 1000x (in 1M HCl) |
| L-Glutamine | 200 | 125x (in H2O) |
| Glycine | 100 | 1000x (in H2O) |
| L-Histidine | 100 | 1000x (in H2O) |
| L-Isoleucine | 100 | 500x (in 1M HCl) |
| L-Leucine | 100 | 500x (in 1M HCl) |
| L-Lysine | 100 | 1000x (in H2O) |
| L-Methionine | 100 | 500x (in 1M HCl) |
| L-Phenylalanine | 100 | 200x (in H2O) |
| L-Proline | 100 | 1000x (in H2O) |
| Hydroxy-L-Proline | 100 | 1000x (in H2O) |
| L-Serine | 100 | 1000x (in H2O) |
| L-Threonine | 200 | 250x (in H2O) |
| L-Tryptophan | 100 | 100x (in H2O) |
| L-Tyrosine | 100 | 250x (in 1M HCl) |
| L-Valine | 100 | 250x (in H2O) |
| p-Aminobenzoic acid (Vitamin B10) | 0.4 | 1000x (in H2O) |
| Biotin (Vitamin B7) | 0.4 | 500x (in H2O) |
| Folic acid (Vitamin B9) | 1.6 | NA |
| Niacinamide (Vitamin B3) | 2 | 1000x (in H2O) |
| β-Nicotinamide adenindinucleotide (NAD) | 5 | 1000x (in H2O) |
| Pantothenate calcium salt (Vitamin B5) | 4 | 1000x (in H2O) |
| Pyridoxal (Vitamin B6) | 2 | 1000x (in H2O) |
| Pyridoxamine dihydrochloride | 2 | 1000x (in H2O) |
| Riboflavin (Vitamin B2) | 4 | NA |
| Thiamine hydrochloride (Vitamin B1) | 2 | 1000x (in H2O) |
| Vitamine B12 | 0.2 | NA |
| Adenine | 40 | NA |
| Guanine | 40 | NA |
| Uracil | 40 | NA |
| CaCl_2_ x 2H_2_O | 10 | 1000x (in H2O) |
| NaC_2_H_3_O_2_ x 3H_2_O | 5,400 | 40x (in H2O) |
| NaHCO_3_ | 5,000 | 40x (in H2O) |
| NaH_2_PO_4_ x H_2_O | 6,400 | 100x (in H2O) |
| Na_2_HPO_4_ x 2H_2_O | 14,700 | NA |
| **Glucose** | **50 mM** | 40x (in H2O) |

**Supplementary Table 3. Reactions predicted to be conditionally essential in all three simulated in vivo conditions.**

|  |  | mean relative growth rate | | | |
| --- | --- | --- | --- | --- | --- |
| reaction ID (BIGG) | **Reaction**  **name** | **rich *in vitro*** | **Nasal-like** | **Plasma-like** | **CSF-like** |
| PRAIS_1 | Phosphoribosylaminoimidazole synthetase | 0.999 | 0.000 | 0.000 | 0.000 |
| GLUPRT | Glutamine phosphoribosyldiphosphate amidotransferase | 0.999 | 0.000 | 0.000 | 0.000 |
| PRFGS | Phosphoribosylformylglycinamidine synthase | 0.999 | 0.000 | 0.000 | 0.000 |
| ADSL1r | Adenylsuccinate lyase | 1.000 | 0.000 | 0.000 | 0.000 |
| ADSS | Adenylosuccinate synthase | 1.000 | 0.000 | 0.000 | 0.000 |
| OMPDC | Orotidine-5-phosphate decarboxylase | 1.000 | 0.000 | 0.000 | 0.000 |
| ORPT | Orotate phosphoribosyltransferase | 1.000 | 0.000 | 0.000 | 0.000 |
| IMPD | IMP dehydrogenase | 1.000 | 0.000 | 0.000 | 0.000 |
| ADSL2r | Adenylosuccinate lyase | 1.000 | 0.000 | 0.000 | 0.000 |
| PRASCSi | Phosphoribosylaminoimidazolesuccinocarboxamide synthase | 1.000 | 0.000 | 0.000 | 0.000 |
| AICART | Phosphoribosylaminoimidazolecarboxamide formyltransferase | 1.000 | 0.000 | 0.000 | 0.000 |
| IMPC | IMP cyclohydrolase | 1.000 | 0.000 | 0.000 | 0.000 |
| ASPCT | Aspartate carbamoyltransferase | 1.000 | 0.002 | 0.003 | 0.002 |
| GMPS2 | GMP synthase | 1.000 | 0.014 | 0.013 | 0.014 |
| GARFT | Phosphoribosylglycinamide formyltransferase | 0.999 | 0.026 | 0.035 | 0.024 |
| PRAGSr | Phosphoribosylglycinamide synthase | 1.000 | 0.059 | 0.076 | 0.058 |
| DHORTS | Dihydroorotase | 1.000 | 0.094 | 0.111 | 0.091 |

### Supplementary Figures

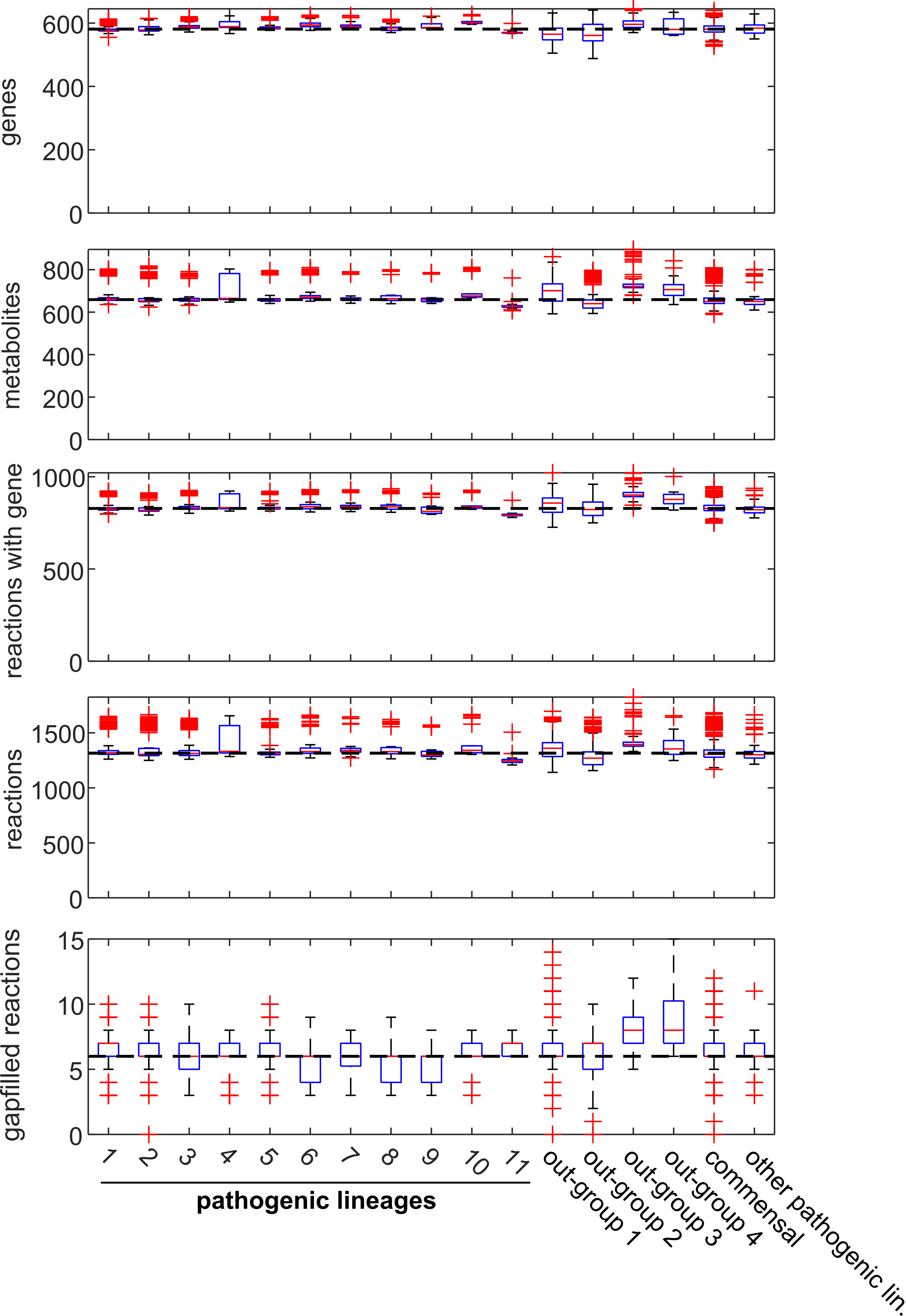

**Supplementary Figure 1. Box plots comparing the number of (from top to bottom) genes, metabolites, gene-associated reactions, reactions, and gapfilled reactions in *S. suis* GSMMs from different lineages.**

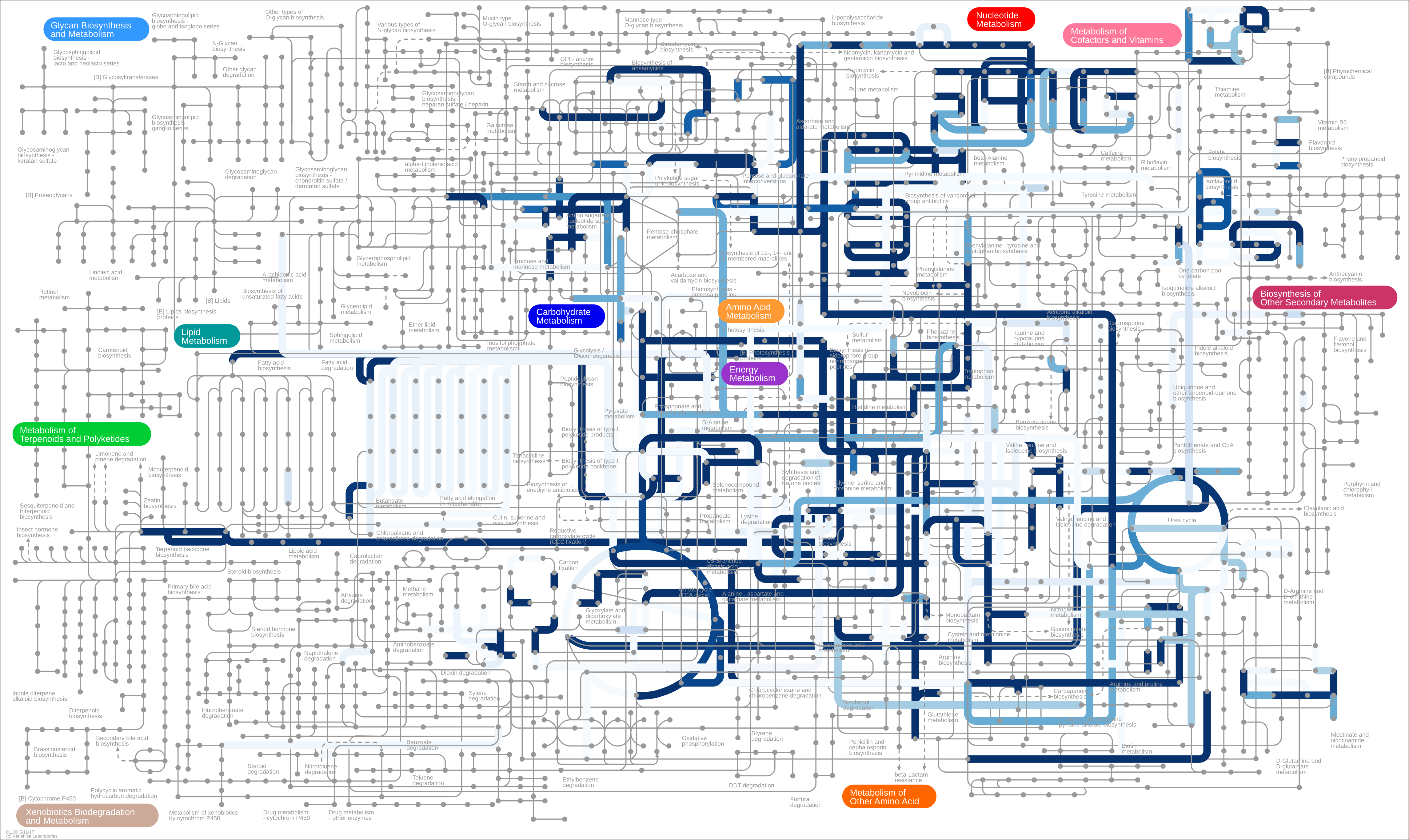

**Supplementary Figure 2. Mapping of gene-associated pan-reactome to the metabolic network.** Mapping was performed by first extracting KEGG reaction IDs for all gene-associated reactions in the pan-reactome (wherever this information was available), and then overlaying these KEGG reaction IDs on a generic metabolic network using iPath3 (*2*) (<https://pathways.embl.de/ipath3.cgi>). Reactions were color-coded by reaction prevalence across models (lowest to highest prevalence: light blue to dark blue).

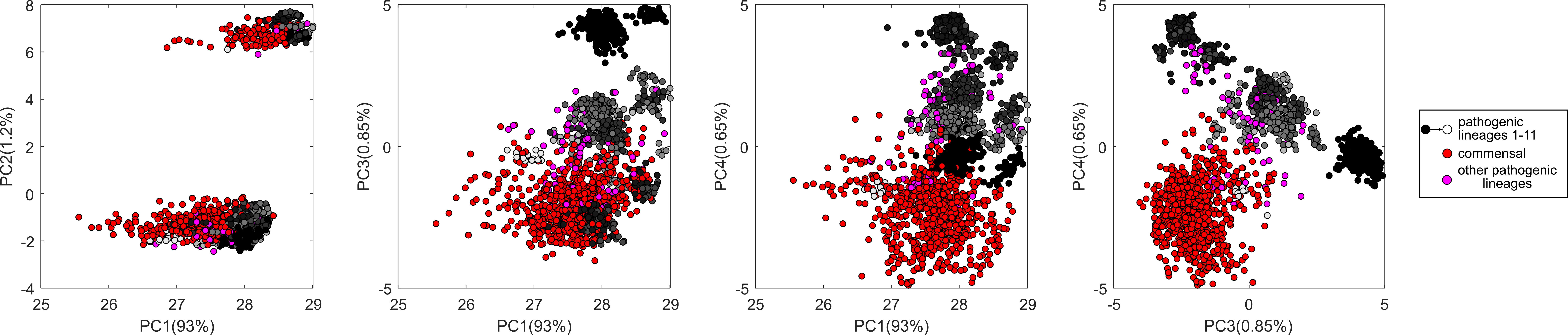

**Supplementary Figure 3. Principal component analysis using the binary matrix of present/absent gene-associated reactions across all models that do not belong to out-groups.** Shown: 2D-plots of principal components 1 to 4. Models were color-coded by lineage as reported in (*1*). % denotes the % of variance explained by the respective principal component.

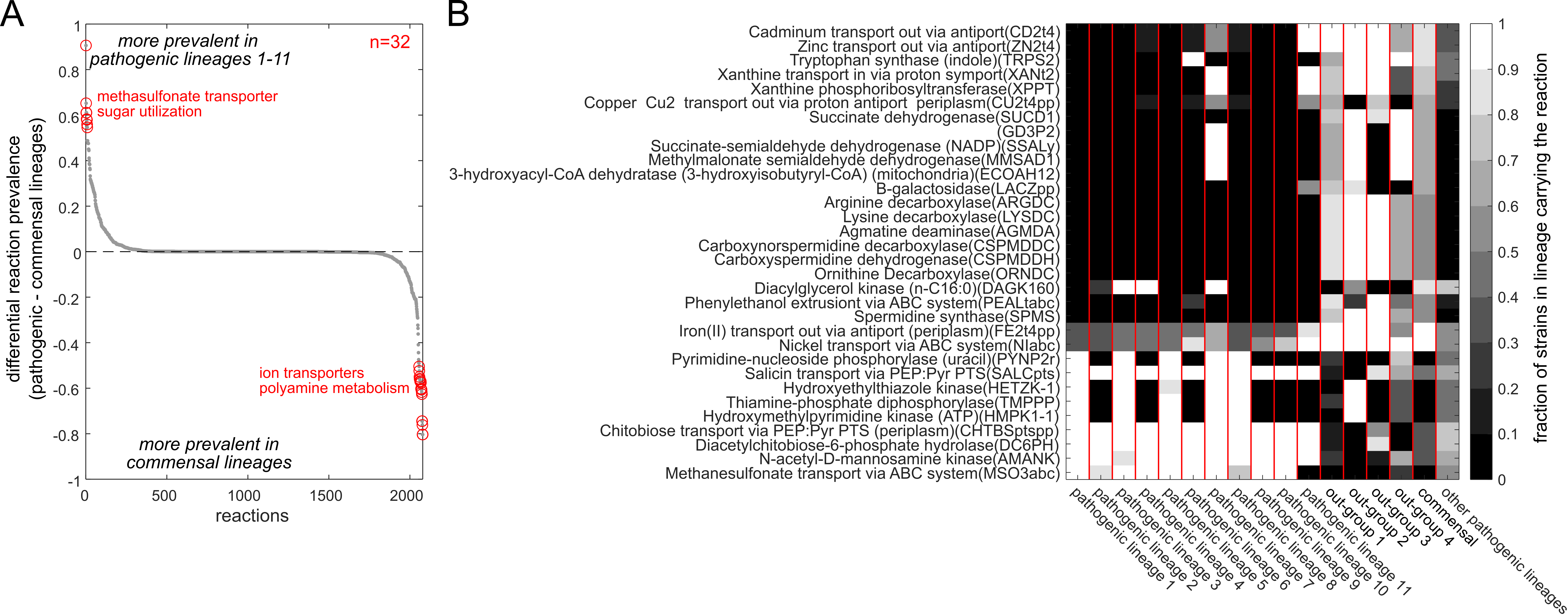

**Supplementary Figure 4. Comparing the prevalence of gene-associated reactions in pathogenic and commensal lineages. A)** Difference in gene-associated reaction prevalence between pathogenic lineages 1-11 and the commensal lineages (lineages defined as reported in (*1*)). Positive values denote higher prevalence in pathogenic lineages, negative values denote higher prevalence in commensal lineages. Highlighted in red: 32 reactions with a difference in prevalence >= 0.5, and some of the pathways they belong to. **B)** Reaction prevalence shown for these 32 reactions with differential reaction prevalence, split by lineages.

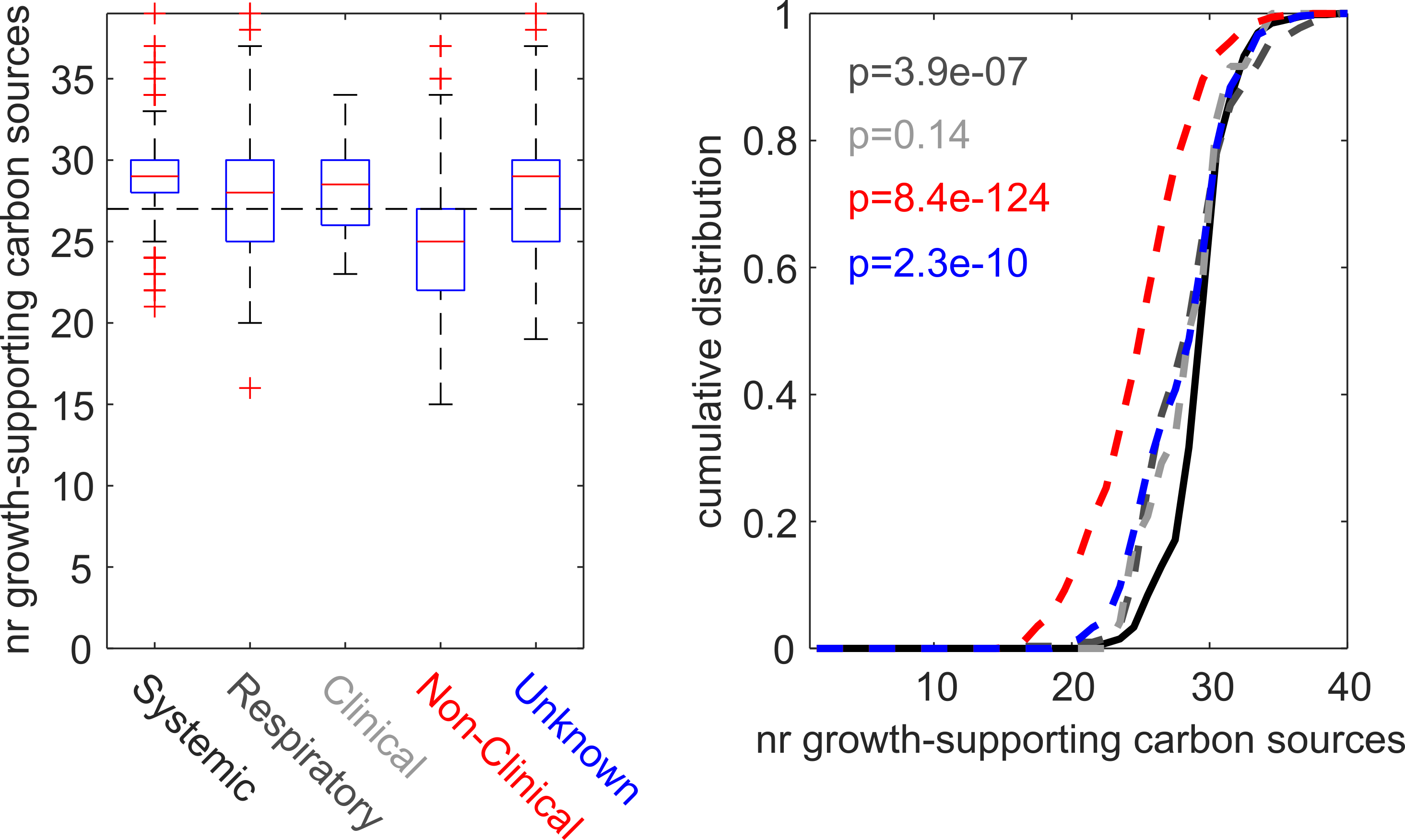

**Supplementary Figure 5. Number of growth-supporting carbon sources across models split by reported disease state.** Left: Boxplots of the number of carbon sources predicted to support growth in each strain, split by disease states, as reported in (*1*). Right: same data but shown as cumulative distributions. Black continuous line: Systemic disease isolates. Dark gray dashed line: respiratory disease isolates. Light gray: clinical isolates. Red dashed line: non-clinical isolates. Blue dashed line: unknown disease state. P-values show the result of a two-sided Wilcoxon ranksum test (testing whether the two sample distributions stem from continuous distributions with equal medians) in reference to systemic isolates.

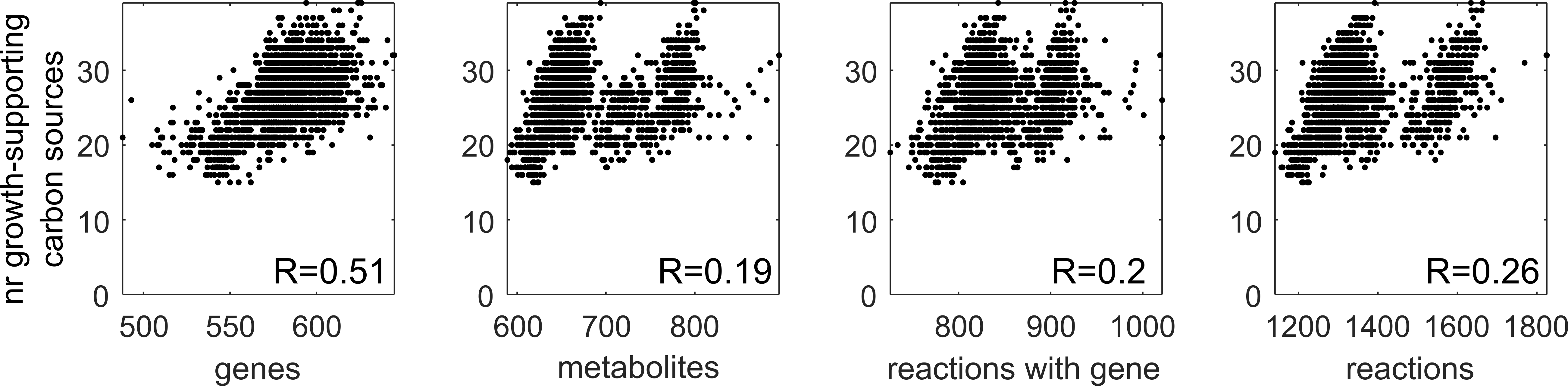

**Supplementary Figure 6. Correlation between the number of growth-supporting carbon-sources (y-axis) and different model features.** From left to right: the number of genes, the number of unique metabolites, the total number of reactions, and the number of reactions having a gene associated, across all models. R denotes the respective Pearson correlation coefficient.

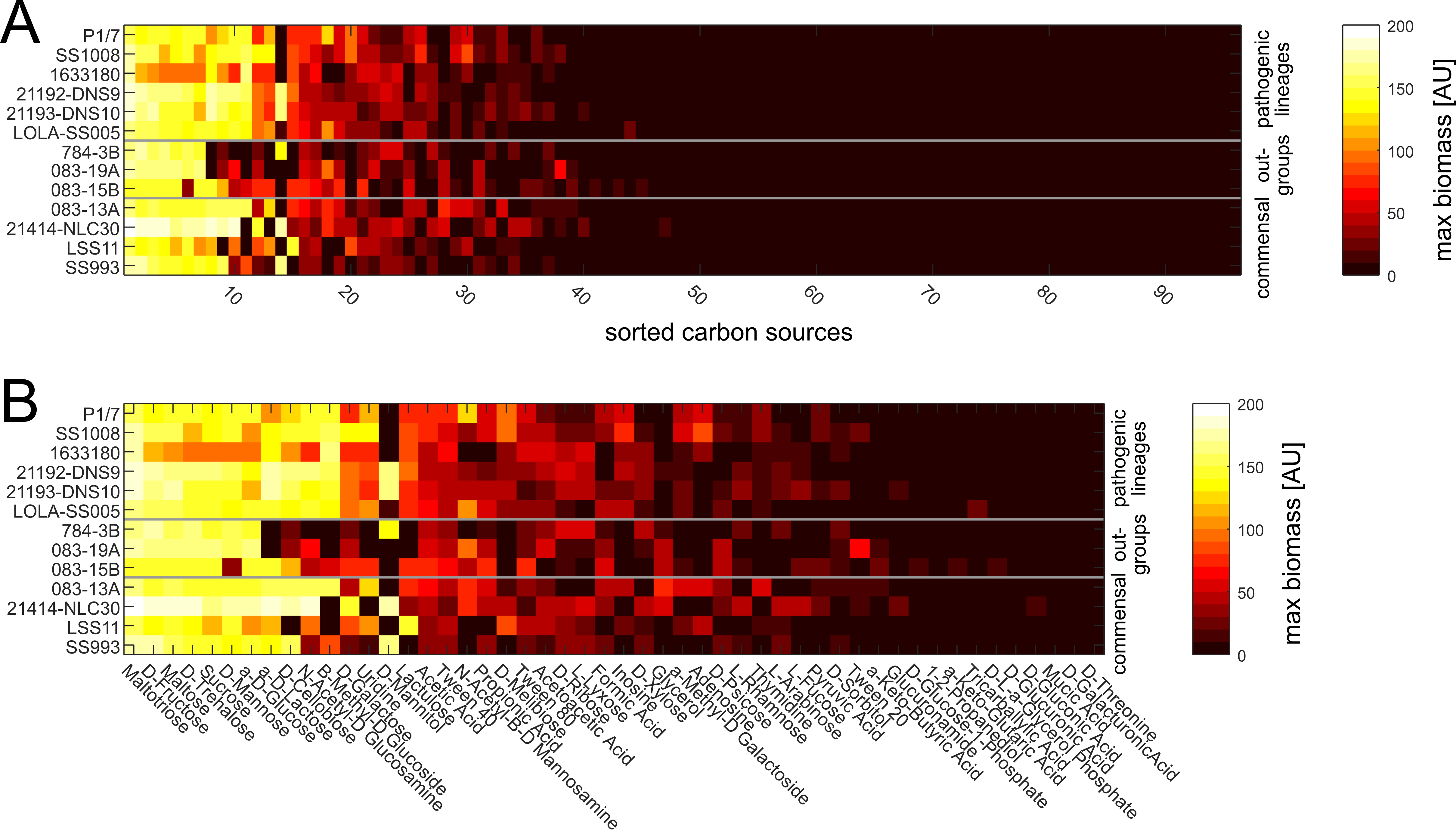

**Supplementary Figure 7. Experimentally determined maximal biomass of *S. suis* strains during growth on 95 individual carbon sources.** Experiments were performed using Biolog plates (i.e. PM1 plates) as described in the main text. Data shown are the mean maximal biomass signals throughout the experiment (3 replicates per strain, 66h duration). **A)** All carbon sources, sorted by their ability to support growth (left to right: highest to lowest biomass production across strains). **B)** Same data, but highlighting the top 50 carbon sources.

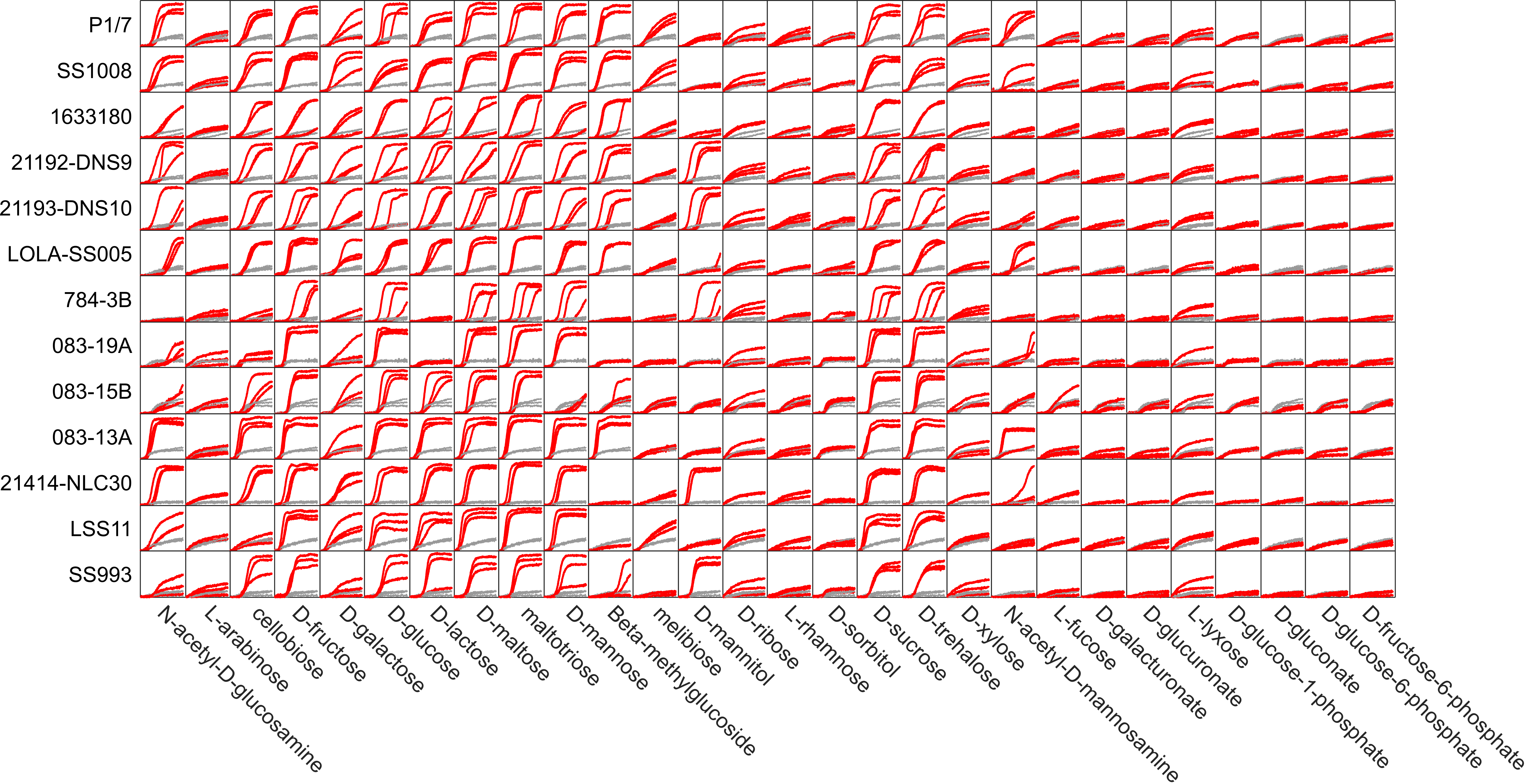

**Supplementary Figure 8. Time courses (using Biolog PM1 plates, see Supplementary Figure 7) of selected *S. suis* strains in 28 carbon sources for which FBA predictions were available.** Red lines: growth in individual carbon sources. Gray lines: signal in negative control (no carbon source). Each line denotes an individual replicate (n = 3 per strain/condition pair). Data were normalized by subtracting the signal at the first time point. X-axis range: 0-70 h. Y-axis range: 0 – 250 AU.

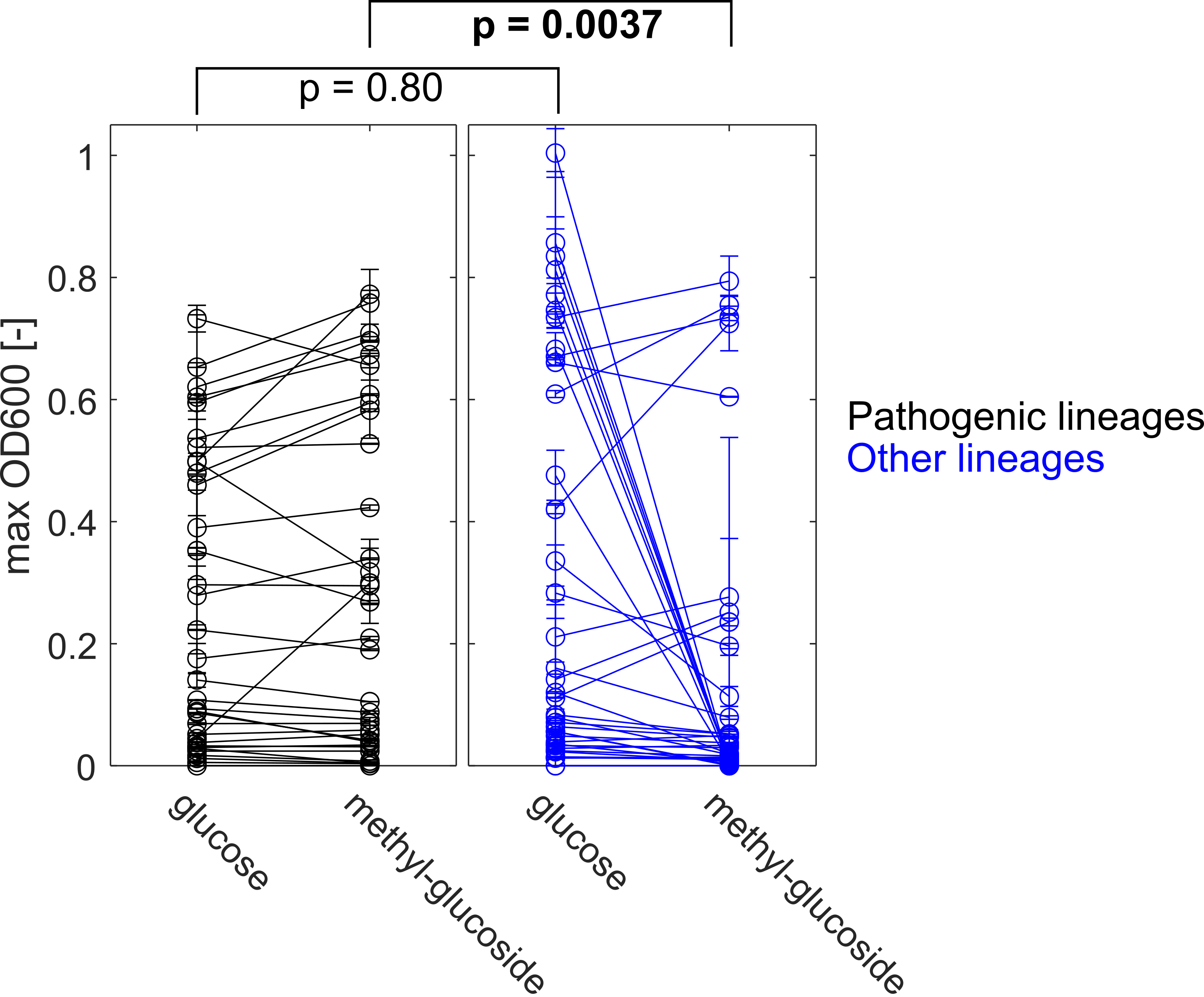

**Supplementary Figure 9. Maximal biomass production of 72 *S. suis* strains grown for 24h in chemically defined medium with glucose or beta-methyl-glucoside as sole carbon sources.** Maximal biomass production was determined as the maximal increase in OD600 within 24h compared to the OD600 signal at the first time point. Left panel (black): strains belonging to pathogenic lineages, as reported in (*1*). Right panel (blue): strains belonging to other (i.e. out-group or commensal) lineages. Lines connect the maximal biomass production of the same strain in each carbon source. Error bars denote standard deviation (n = 2-9).

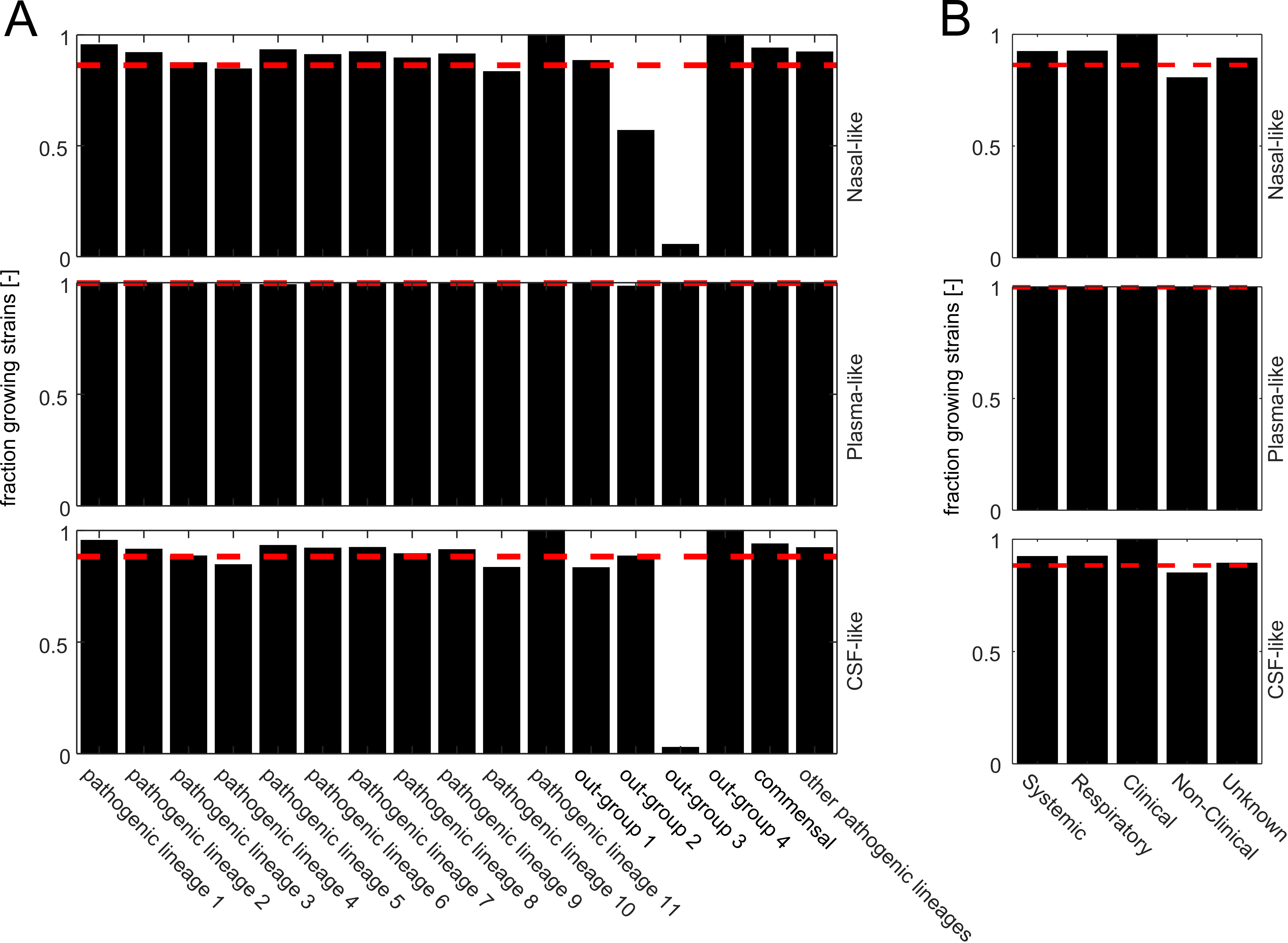

**Supplementary Figure 10. Fraction of strains predicted to grow in inferred *in vivo* environments split by lineage (A), or by disease state (B).** Strains were predicted to grow if their relative growth rate (compared to the growth rate in simulated *in vitro* rich medium) is larger than 0.001. Red dashed line: total fraction of strains predicted to grow in inferred in the respective vivo environment. Strains were categorized by lineage and disease state as reported in (*1*).
